# Combination of *kdr* mutation and detoxification gene expression associated with high tolerance for permethrin in a resistant *Aedes aegypti* population

**DOI:** 10.1101/2020.06.22.164483

**Authors:** Tse-Yu Chen, Chelsea T. Smartt, Dongyoung Shin

**Author notes:** Corresponding Authors (DS), (CTS).

## Abstract

*Aedes aegypti*, as one of the vectors transmitting several arboviruses, is a main target in mosquito control programs. Permethrin remains the major adulticide used to control these mosquitoes. The increasing percentage of permethrin resistant *Aedes aegypti* has become an important issue around the world. Knockdown resistance (*kdr*) is one of the major mechanisms related to permethrin resistance. On the other hand, detoxification genes including cytochrome P450 monooxygenases (P450) and glutathione S-transferases (GSTs) are also suggested as permethrin resistance apparatus. Here we selected a permethrin resistant (p-s) *Aedes aegypti* population from Florida and compared its mortality after exposure, median lethal dose (LD50), adult survivorship and larval development to several field populations. We used allele-specific PCR genotyping of the S989P, V1016I and F1534C sites in the sodium channel gene and gene expression analysis of several p450 and GSTs genes before and after permethrin exposure to determine their involvement in permethrin sensitivity between *Ae. aegypti* populations. Results indicated the p-s population had the highest resistance to permethrin based on LD50 and the mortality test. The larval development time did not significantly differ between the populations, however the p-s adults survived longer than the other populations. In the genotype study, p-s population had mostly homozygous mutations in all three mutant sites of the sodium channel gene. Detoxification gene expression studies showed that two p450 genes, AAEL009124 (*CYP6N12*) and, AAEL009127 (*CYP6M11*), were upregulated and, accession # AAEL006802, AAEL014891 (*CYP6P12*) and AAEL014619 (*CYP9J22*) were downregulated after 120 minutes of permethrin exposure in the p-s population. These results suggest that in highly permethrin resistant *Aedes aegypti* populations both *kdr* mutations and xenobiotic metabolism might be involved. Involvement of multiple mechanisms to achieve resistance to permethrin supports the need for implementing comprehensive mosquito control measures, such as an integrated pest management strategy, so that selection pressure for resistance is decreased without compromising control efforts while new methodologies are being developed.

**Author summary:** Pyrethroids have been applied as a major type of insecticide targeted at *Aedes aegypti*, a key vector in the transmission of several flaviviruses. Resistance to pyrethroids has emerged and has become a worldwide threat to mosquito control. Pyrethroid resistance usually occurs with knockdown resistance (*kdr*) where the voltage gated sodium channel is mutated. We selected a permethrin resistant (p-s) *Aedes aegypti* population from Florida and, along with two other field populations, examined three mutation sites, S989P, V1016I and F1534C. The data showed the p-s population had the most homozygous mutations which correlated to the permethrin resistance level. Besides *kdr*, detoxification genes also have been identified to have pyrethroid metabolizing abilities. We found two cytochrome P450 monooxygenases genes, *CYP6N12* and *CYP6M11*, were overexpressed in the p-s population after permethrin exposure, suggesting a role in resistance to permethrin. Together, our results provide information about potential mechanisms used in major vectors with high insecticide resistance.

## Introduction

Mosquitoes are the most common vectors of parasitic and viral diseases. Mosquito-borne diseases have a tremendous human impact, resulting in over one million deaths every year. The *Aedes aegypti* mosquito is the primary vector of many arboviruses, including yellow fever, dengue, chikungunya, and zika viruses [1,2]. The application of insecticides is a widely used approach to control mosquitoes; however, insecticide use leads to the development of insecticide resistance within mosquito populations and also has detrimental effects on beneficial organisms [3]. Therefore, there continues to be a need to understand the mechanisms that are involved in development of insecticide resistance in order to mitigate resistance so that traditional control methods can still be used while new non pesticide control strategies are being developed.

Mechanisms of insecticide resistance in mosquitoes include behavioral changes [4], alterations of the cuticle to reduce insecticide penetration [5], increased detoxification metabolism [6,7], and changes to target sites on sodium channel proteins [8]. Pyrethroids and DDT target the voltage-gated sodium channel of insect neurons. Single amino acid substitutions in the sodium channel have been associated with resistance to those insecticides [9]. This form of resistance, known as knockdown resistance (*kdr*), has been observed in several insect species including *Ae. aegypti* [10]. In many insect species, mutations related to both pyrethroid and DDT resistances have mainly been located in the IIS6 region of the voltage-gated sodium channel gene. The replacement of a Leucine for a Phenylalanine at amino acid 1014 (Leu1014Phe/Ser) was the most common [11,12]. In *Ae. aegypti*, a number of *kdr* mutations have been identified (S989P, I1011M/V, V1016G/I, F1269C, F1534C). At least two of these are known to be related to pyrethroid resistance: 1016 (Val to Ile or Gly) and 1534 (Phe to Cys) in the IIS6 and IIIS6 segments of the voltage-sensitive sodium channel gene, respectively [10,13–15].

The detoxification of insecticides in mosquitoes involves three major metabolic detoxification gene families: cytochrome P450s (P450s), esterases, and glutathione S-transferases (GSTs). P450s are one of the largest gene families in all living organisms and are known to respond to pyrethroid insecticide; esterase is more active against organophosphate [16]. GSTs are soluble dimeric proteins that are involved in the metabolism, detoxification, and excretion of a large number of endogenous and exogenous compounds [17,18]. The alteration of gene expression of insect P450s and GSTs results in increased enzymatic activities. Consequently, this alteration enhances the metabolic detoxification of insecticides and the development of insecticide resistance [6]. The increased gene expression levels in P450s has also been reported to be involved in the development of insecticide resistance [19].

Insecticide resistance is energetically costly and results in changes to the physiology and behavior of mosquitoes, which in turn may alter the vector competence. There are many reports showing that changes in vector competence correlate positively or negatively with changes in pathogen transmission and other biological factors affecting vectorial capacity [20–22]. Development, life span, and fecundity are the biological factors with the strongest impact on vectorial capacity in *Ae. aegypti* and these factors are altered in insecticide resistant populations [23,24]. Although chemical control remains the best method for controlling mosquitoes and the pathogens they transmit, insecticide-resistant mosquitoes can increase the risk of mosquito-borne disease transmission. This would greatly hinder mosquito control in its ability to effectively decrease the number of mosquito pests. Determination of insecticide resistance mechanisms in Florida mosquito populations will provide knowledge that can be used for mosquito control. The impact of insecticide resistance on mosquito biology—especially relating to mechanisms of vector competence—can inform programs that operate to mitigate the impact of re-emerging mosquito-borne pathogens and the increase in insecticide resistance.

## Methods

### Mosquitoes

The *Aedes aegypti* larvae was collected in Key West, Florida (Key West population) and Vero Beach, Florida (Vero population). Field-collected mosquitoes were sorted by species and maintained in the laboratory under standard insectary conditions (26 °C, 14 h/10 h light/dark period, 70% relative humidity). Mosquito colonies are maintained by feeding female mosquitoes on blood from chickens following approved standard protocols (IACUC protocol 201807682) and eggs are collected [25]. Generations F_13_ to F_20_ were used in this study.

### Selection for permethrin resistance in *Ae. aegypti* and mortality assessment

Adult Key West *Ae. aegypti* mosquitoes were separated in males and females and exposed to permethrin using a bottle bioassay, respectively [26–28]. The survivors were allowed to mate and blood fed to generate the population (p-s population). Thus, we exposed successive generations of mosquitoes to increasing concentrations of permethrin so that each generation suffered ∼30% mortality from the exposure. We used adult mosquitoes of the F_13_ to F_20_ generation (p-s population) to compare with other field populations collected at the same time with the p-s population. One of the populations, Key West is the parental population of the p-s population.

In the mortality experiment, female and males were placed separately in 1ml bottles coated with equivalent amounts of 43 μg permethrin and the mortality counted after 90 minutes [27].

In the Median Lethal Dose (LD50) study, p-s population, Key West and Vero populations were used. Around 20 mosquitoes were exposed to different concentrations of permethrin in coated 250 ml bottles. The mosquito knockdown percentage was recorded after 10 minutes exposure and the permethrin concentration adjusted so that the final concentration yielded around 50% knockdown for each population.

### Assessment of phenotypic insecticide resistance in selected *Ae. aegypti*

For genomic DNA extraction, single mosquito bodies from the p-s population, Key West and Vero populations were collected in 1.5ml tubes containing 100ul of 25 mM NaCl, 10 mM Tris-Cl pH 8.0, 1 mM EDTA, 200 μg/ml proteinase K and homogenized using a pestle. After incubating the mixture for 30 min at 37 °C and inactivation of proteins with proteinase K at 95 °C for 5 min, the mixture was centrifuged at 12,000xg for 5 minutes, and the supernatant containing DNA used for PCR. The sodium channel gene amplification by PCR was used to detect S989P, V1016I, and F1534C mutations in each of the *Ae. aegypti* populations.

PCR was conducted to check individual mosquito *kdr* mutation sites. Two reactions were assembled the same except that one contained a susceptible-specific primer and the other contained a mutant-specific primer [29](S1 Table). A PCR diagnostic test was performed in accordance with the standard procedure with a total reaction volume of 25 μl, consisting of 100–200 ng genomic DNA, 2.5 μl 10x buffer, 0.5 mM dNTP mix and 2.5 U JumpStart AccuTaq LA DNA Polymerase Mix. PCR conditions were one cycle of 96 °C for 30 s, then 35 cycles of 94 °C for 15s, 55 or 60 °C (60 °C for V1016G and F1534C, 55 °C for S989P) for 30 s, and 68 °C for 1 min, followed by one cycle of 68 °C for 5 min. PCR products were checked by electrophoresis on 1.5% agarose gels in TEB buffer. Bands were visualized by GelRed® staining. The size of the PCR products for the detection of *kdr* alleles were 240 bp (S989P), 348 bp (V1016I) and 284 bp (F1534C), whereas the size of the products used as allele-nonspecific outer primers were 594 bp (S989P), 592 bp (V1016I), and 517 bp (F1534C).

### Comparison analysis of the survivorship and development between permethrin-resistant and field populations

The survivorship was compared between the p-s population with two field populations (Vero and Key West) and one lab population of *Ae. aegypti* (Rockefeller). Adult mosquitoes that emerged on the same day were collected and used in this experiment. Twenty-five female adults from each population were placed in separate 8oz soup cups covered with mesh for a total of 10 replicates and provided 20% sucrose water. Cups were placed in standard insectary conditions (26 °C, 14 h/10 h light/dark period, 70% relative humidity). The mortality was recorded each day, until no more mosquitoes survived. Tukey’ s multiple comparisons test was used to analyze each population’ s survivorship.

Larval development period was also compared between the p-s population and three other populations of *Ae. aegypti* (Vero, Key West and Rockefeller). In order to hatch the eggs simultaneously, a piece of egg paper was placed in a flask containing water and connected to a vacuum to remove the air. After 30 minutes, 25 1st instar larvae were placed in a plastic cup containing 150ml water and 0.15-gram larvae food (brewer yeast and albumin mixture=1:6). Each population had 10 replicated cups and were placed in the incubator (14:10 light cycle and 26 °C). Both the status of larvae instar and mortality were recorded every day. Once pupae were observed, the pupae were removed from the cup to a falcon tube and the sex recorded after emergence. Tukey’ s multiple comparisons test was used to analysis each population’ s larval development time.

### Comparison of gene expression levels in detoxification genes using qRT-PCR

To test the detoxification gene expression level, the p-s population and the Key West population were used. Five-day-old female mosquitoes were tested in 250 ml bottles coated with 15ug of permethrin. Thirty of the p-s population exposed to permethrin for 30 minutes and 120 minutes were collected, as well as 30 unexposed mosquitoes from the same population. All of the Key West population were dead after 120 minutes of exposure to permethrin; thus, sampling from the Key West population was from the unexposed group and those exposed for 30 minutes. Total RNA from ten female mosquitoes per population per group was extracted using Trizol to determine the targeted gene expression [30]. The primers specific to the genes of interest (S2 Table) were designed to determine changes in detoxification gene expression by quantitative Real-Time PCR (qRT-PCR). The related gene expression level was normalized to the expression of the *Ae. aegypti* ribosomal protein S7 gene (GenBank Accession # AY380336) [25]. The expression level was compared with non-permethrin exposed mosquitoes. The gene expression in each group was compared by delta-delta Ct value analysis. Standard deviation was calculated. Wilcoxon Method was used to calculate p value and only p<0.05 was counted as significantly different.

## Results

### Mortality and LD50 for permethrin between permethrin-resistant and field populations

Resistance level of each mosquito population to permethrin was assessed and compared for the p-s population, two field populations from Florida (Key West and Vero) and the susceptible Rockefeller population using the standard CDC bottle bioassay. Mortality for the p-s population was 6-fold lower than the Rockefeller population and the field populations at the new CDC diagnostic dose (43 ug/bottle) (Table 1)[27].

**Table 1.** Mortality of *Ae. aegypti* populations following exposure to permethrin.

| *Ae. aegypti* population | No. tested | % Mortality* |
| --- | --- | --- |
| Rockefeller | 200 | 0.983±0.005 |
| Vero | 200 | 0.980±0.016 |
| Key West | 200 | 0.815±0.087 |
| p-s | 420 | 0.235±0.009 |

\*Values are the mean ± SE.

Permethrin median lethal dose was tested in Vero, Key West and p-s populations. The median lethal dose (LD50) for the Vero population was 100μg, for Key West the dose was 240μg and the p-s population was 5mg (Table 2).

**Table 2.** Toxicity of permethrin to the *Ae. aegypti* populations.

| *Ae. aegypti* population | No. tested | LD50* |
| --- | --- | --- |
| Vero | 96 | 100µg |
| Key West | 80 | 240µg |
| p-s | 45 | 5000ug |

\*Lethal dose required to kill 50% of the population in ug. A minimum of three assays were performed per population.

### Analysis of survivorship and larval development of permethrin-resistant versus field populations

The adult survivorship of the selected population (p-s), two field *Ae. aegypti* populations (Key West and Vero) and susceptible Rockefeller population was assessed to determine the effect of permethrin resistance on survivorship, a major component of vectorial capacity. The p-s population lived significantly longer than the parental population, Key West. However, the p-s population total survival time did not have significant differences to Vero and Rockefeller populations (Table 3).

**Table 3.** Analysis of survivorship of the *Ae. aegypti* populations.

| Population | Total survival time |
| --- | --- |
| p-s | 17.96<sup>a</sup> |
| Key West | 11.80<sup>b</sup> |
| Rockefeller | 16.19<sup>ab</sup> |
| Vero | 17.47<sup>ab</sup> |

<sup>ab</sup> Levels not connected by the same letter are significantly different.

In the larval development study using the same populations, the Rockefeller population took the shortest time for pupation but both male and female development time was not significantly different between different populations (Table 4).

**Table 4.**
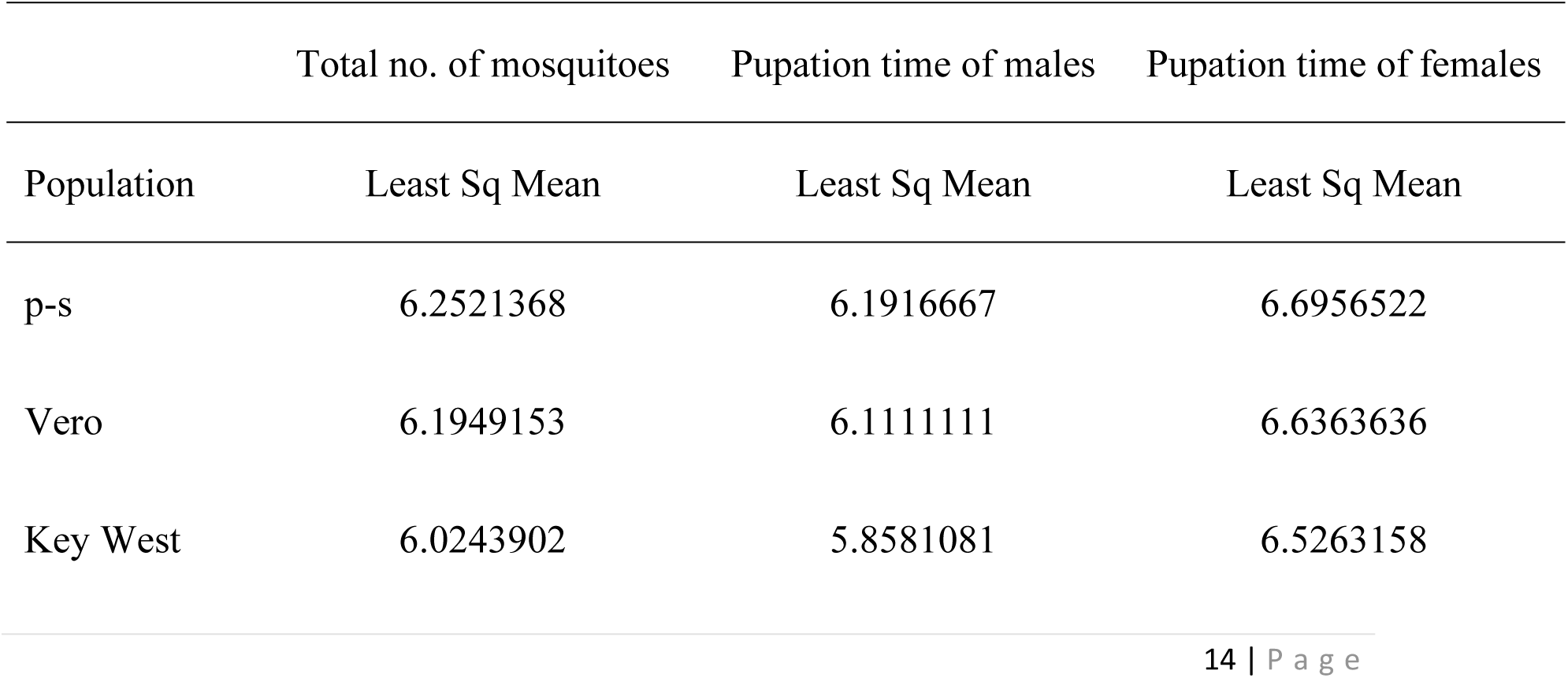
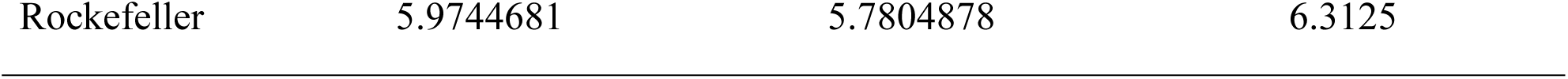
Analysis of larval developmental time for the *Ae. aegypti* populations.

### Assessment of the kdr mutations responsible for phenotypic insecticide resistance in three *Ae. aegypti* populations

Three mutations were identified from the genomic DNA from both female and male *Ae. aegypti*: S989P (TCC-CCC), V1016I (GTA-ATA), and F1534C (TTC-TGC) in a subset of each population (Table 5). To further investigate the point mutation status, allele-specific primers per each mutation site on the *kdr* gene were used to screen the mutation sites in the three mosquito populations (p-s, Vero and Key West) (Table 5). The allele-specific PCR detected the mutation in all three mutant sites, S989P, V1016I and F1534C. The p-s population had 100% mutation in F1534C, 97% mutation in V1016I and 74% mutation in V989P. The Vero population had 77% mutation in F1534C, 67% mutation in V1016I and 56% mutation in V989P. The Key West population had 100% mutation in F1534C, 64% mutation in V1016I and 87% mutation in V989P (Table 5). Moreover, the population from Key West showed the most individuals as heterozygous (80%), and the p-s population showed mostly homozygous mutations (53%) for the mutation sites tested.

**Table 5.** Percentage of mutations in the sodium channel gene in populations of.

| F1534C |  | Number* |  |  | Percentage (%) |  |  |
| --- | --- | --- | --- | --- | --- | --- | --- |
| Population | Gender | F | F+C | C | F | F+C | C |
| p-s | Male | 0 | 6 | 4 | 0.00 | 60.00 | 40.00 |
|  | Female | 0 | 13 | 16 | 0.00 | 44.83 | 55.17 |
| Vero | Male | 1 | 9 | 0 | 10.00 | 90.00 | 0.00 |
|  | Female | 8 | 18 | 3 | 27.59 | 62.07 | 10.34 |
| Key West | Male | 0 | 10 | 0 | 0.00 | 100.00 | 0.00 |
|  | Female | 0 | 25 | 4 | 0.00 | 86.21 | 13.79 |
| V1016I |  | Number* |  |  | Percentage (%) |  |  |
| Population | Gender | V | V+I | I | V | V+I | I |
| p-s | Male | 0 | 5 | 5 | 0.00 | 50.00 | 50.00 |
|  | Female | 1 | 4 | 24 | 3.45 | 13.79 | 82.76 |
| Vero | Male | 1 | 9 | 0 | 10.00 | 90.00 | 0.00 |
|  | Female | 12 | 16 | 1 | 41.38 | 55.17 | 3.45 |
| Key West | Male | 1 | 9 | 0 | 10.00 | 90.00 | 0.00 |
|  | Female | 13 | 16 | 0 | 44.83 | 55.17 | 0.00 |
| S989P |  | Number* |  |  | Percentage (%) |  |  |
| Population | Gender | S | S+P | P | S | S+P | P |
| p-s | Male | 3 | 6 | 1 | 30.00 | 60.00 | 10.00 |
|  | Female | 7 | 10 | 12 | 24.14 | 34.48 | 41.38 |
| Vero | Male | 3 | 6 | 1 | 30.00 | 60.00 | 10.00 |
|  | Female | 14 | 10 | 5 | 48.28 | 34.48 | 17.24 |
| Key West | Male | 1 | 9 | 0 | 10.00 | 90.00 | 0.00 |
|  | Female | 4 | 25 | 0 | 13.79 | 86.21 | 0.00 |

\*Column numbers indicate the total number of mosquitoes tested.

### Comparison of gene expression levels for detoxification genes in three *Ae. Aegypti* populations

Based on the literature and the previous RNAseq data analysis with the p-s population (data not shown), several cytochrome P450 monooxygenases (P450) and glutathione S-transferase (GST) genes, which are detoxification genes for pyrethroids, were selected [13,31](S2 Table) and their gene expression assessed among the mosquito populations used in this study (Figure 1).

**Figure 1.**
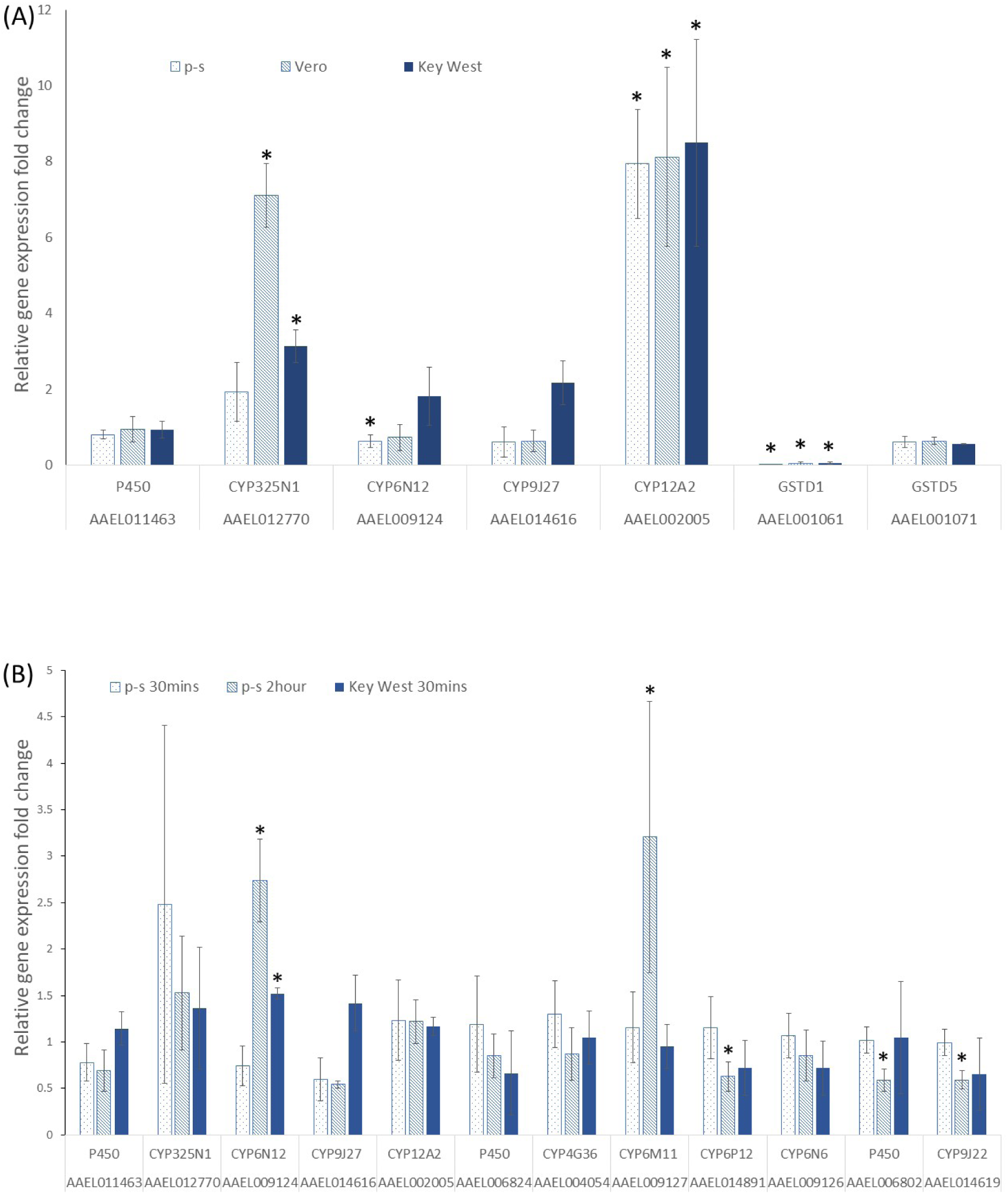
Comparison of detoxification gene expression among populations. (A) Relative detoxification gene expression between p-s to Rock (Rockefeller strain), Vero to Rock and Key West to Rock without permethrin treatment. (B) Cytochrome p450 gene expression after permethrin treatment in the p-s population and the parental population (Key West) compared to non-treatment group. Asterisks (*) indicate a significant difference by Wilcoxon Method (p<0.05).

Prior to exposure to permethrin: The Rockefeller strain was used to assess baseline detoxification gene expression levels and expression compared with the p-s and field populations. One of the GST genes, *GSTD1* (AAEL001061), was significantly downregulated without permethrin exposure in p-s population and field populations compared to the lab population but not the other GST gene *GSTD5* (AAEL001071) (p value > 0.05). The *CYP12A2* (AAEL002005) showed highly elevated expression levels in all the field populations and in the p-s population without insecticide exposure compared to the lab population. The p-s population did not show elevated gene expression in most of the detoxification genes except for *CYP12A2*. The *CYP325N1* (AAEL012770) was upregulated in the filed populations (Vero and Key West) but not p-s population. The *CYP6N12* (AAEL009124) was downregulated in the p-s population compared to all other tested populations (Figure 1A).

Post exposure to permethrin: Several P450 genes were selected and the gene expression level of those genes were compared in the same populations without permethrin treatment. The *CYP6N12* expression level was significantly increased in the p-s population after 120 min of permethrin treatment while the Key West population showed in increase in expression of this gene after 30 minutes (Figure. 1B). The *CYP6M11* (AAEL009127) was also highly expressed in the p-s population after 120 min of permethrin treatment and expression decreased for AAEL006802, *CYP6P12* (AAEL014891) and *CYP9J22* (AAEL014619). The other seven P450 genes did not have expression differences after exposure, including AAEL011463, *CYP325N1* (AAEL012770), *CYP9J27* (AAEL014616), *CYP12A2* (AAEL002005), AAEL006824, *CYP4G36* (AAEL004054) and *CYP6N6* (AAEL009126).

Expression levels of GST genes, *GSTD1* (AAEL001061), *GSTD5* (AAEL001071), *GSTE6* (AAEL007946), *GSTE2* (AAEL007951) and *GSTE5* (AAEL007964), did not change in the p-s population and field populations after permethrin exposure (S1 Figure). Two cuticle-related genes (AAEL011897 and AAEL017542) were tested following permethrin treatment to determine whether permethrin exposure stimulates cuticle thickening. There were no differences in expression of the cuticle genes in any of the populations (data not shown).

## Discussion

Resistance to permethrin in *Aedes aegypti* has been a worldwide issue for mosquito control [3,32]. In this study, we performed a comprehensive screen with a laboratory selected permethrin-resistant *Ae. aegypti* population and evaluated the potential mechanisms that might have contributed to the high resistance.

The p-s population selected from the Key West population had higher resistance compared to the other wild and lab populations (Table 1). The permethrin LD50 study verified that the p-s population had significantly higher dose tolerance for permethrin (Table 2) and corroborates that this population is resistant. The field populations (Vero and Key West) showed higher resistance levels to permethrin compared to the susceptible lab population (Rockefeller).

The elevated insecticide resistance seemed to affect adult longevity of the p-s population (Table 3) as the p-s population had a longer life span than the parental Key West population. These findings are contrary to what was shown in previous studies where pyrethroid resistance in mosquitoes was shown to have negative effects on life-history traits including decreased adult longevity [24,33]. Additionally, longevity is one of the important factors in the measurement of vectorial capacity [34] because it allows infected mosquitoes to transmit the pathogenic virus for a longer period of time. Longer life span and higher tolerance for permethrin could increase potential as a vector for pathogenic virus. In fact, our previous studies using this population showed higher infection, dissemination and transmission rates for dengue virus (Shin, D., personal communication) and higher zika virus replication rate [35].

Insecticide resistance can generally be conferred by increased detoxification metabolism [36,37]. Some of this increased metabolism is initiated by insecticide contact rather than the result of an inherently high metabolism [13]. The p-s population might initiate the detoxification metabolism once exposed to permethrin to decrease the cost to fitness. Furthermore, the p-s population was selected only with permethrin while the parental population, Key West, was likely exposed to different types of insecticide in the wild including permethrin, malathion and naled [38]. The Key West population had significantly lower survivorship than the p-s population perhaps due to the high cost of detoxifying different insecticides. The Vero population, on the other hand, would have been exposed mainly to permethrin in the wild [39] and had no differences to the p-s population in adult longevity.

In the larvae development time experiment (Table 4), both the male development time and the female development time were not significantly different across different populations. Resistance to permethrin was shown to not impact pupation in select isofemale lines from a DDT- and permethrin-resistant *Ae. aegypti* population originally collected in Thailand [40]. However, our data suggested p-s population required longer development time than other field populations (Table 4). This delayed development time in an insecticide-resistant population is consistent with what has previously been seen in laboratory studies [41] and may be caused by the allocation of energy to detoxify the insecticide and a depletion of nutrients required for development [24].

It was previously shown that *Ae. aegypti* in Florida had different levels of resistance to permethrin and *kdr* mutations, and substitutions at positions 1016 and 1534 were the most common [42]. Here we examined 3 *kdr* mutant sites in a permethrin selected *Ae. aegypti* population and 2 field populations and found the p-s population had the highest homozygous mutation rate in the three voltage gated sodium channel gene sites, S989P, V1016I and F1534C. The homozygous mutation of V1016I and F1534C resulted in higher resistance to the pyrethroid deltamethrin than heterozygous mutants [43]. Moreover, the median lethal time (LT50) confers the triple mutated homozygote populations, S989P, V1016G and F1534C, survived longer than triple mutated heterozygote populations under permethrin exposure [44]. Therefore, the high percentage of homozygous mutations for S989P, V1016I and F1534C sites might have contributed to higher resistance to permethrin in the p-s population than the other field populations (Table 2).

Most of the p-s population had mutations in all three sites (71.8%), regardless of presence of homozygous or heterozygous mutations, but only 56.4% of these mutations were found in the Key West population and 28.2% in the Vero population (data not shown). The higher mutation rate reflected a positive correlation with the LD50 to permethrin for the p-s population (Table 2). Although S989P mutation often coexist with V1016I in field permethrin resistant *Ae. aegypti* and presence of a combination of S989P and V1016I alleles is not known to increase resistance levels compared to having only a single mutation [45], the presence of an F1534C mutation is known to reduce the channel sensitivity to permethrin [46] and the combination of V1016I and F1534C alleles have more resistance to the pyrethroid deltamethrin than only V1016I mutation [43]. This is indicative that the p-s population has high resistance to permethrin due to having all three mutant alleles S989P/V1016I/F1534C.

The cytochrome P450 monooxygenases (P450) have been proven to function in xenobiotic metabolism including insecticide resistance in insects [6]. Additionally, several P450 genes were induced after permethrin treatment in *Aedes aegypti* [47,48] and then identified as being capable of pyrethroid metabolism [49]. In our study, *CYP12A2* (AAEL002005) was the only P450 with increased expression in unexposed p-s population and the other wild populations compared to Rockefeller population (Figure 1A). The results corroborate what was shown by Goidin et al. (2017) who showed increased expression of *CYP12A2* in insecticide resistant field mosquitoes [50]. The P450s might not be induced without permethrin exposure therefore decreasing the energy required to maintain functions involved in longevity (Table 3). However, the gene expression of *CYP12A2* did not change after permethrin exposure in any population (Figure 1B). The expression of this gene may be elevated for general defense, including from xenobiotics like pesticides. Two P450 genes, *CYP6N12* (AAEL009124) and *CYP6M11* (AAEL009127), were overexpressed after 2 hours of permethrin exposure in the p-s population compared to non-exposed p-s population (Figure 1B). In a previous study, *CYP6N12* was not induced after 72 hours exposure to permethrin in the larval stage [47,51]. However, the P450 genes had different expression levels between different life stages [52] and different populations [53]. In another study, *CYP6M11* was induced after 48 hours exposure to permethrin in the larval stage [54] and, in addition to results from this study, indicates that *CYP6M11* might be one of the P450 enzymes that detoxify permethrin in *Ae. aegypti* larvae and adults.

Glutathione S-transferase (GST) gene expression and enzymatic activity have been suggested to play an important role in resistance to permethrin in *Ae. aegypti* [7]. Expression of GST genes also showed correlations with larval development, aging [55] and pathogen presence, including dengue and West Nile viruses [56,57], thus alterations in expression may not indicate involvement in resistance. Our data showed that one GST encoding gene, *GSTD1* (AAEL001061), had significantly reduced expression in unexposed p-s, Key West and Vero populations compared to the Rockefeller population (Figure 1A). However, after permethrin exposure, there were no significant differences in GST gene expression compared to exposure groups (S1 Figure). This result suggests that GST gene transcription is not enhanced in these populations of permethrin-resistant *Ae. aegypti*. However, involvement of GST proteins in resistance still needs to be assessed although assignment of an actual role of GST in permethrin resistance might not be straightforward [58,59]. It is critical to investigate permethrin-specific detoxification genes in mosquito populations with variable resistance ratios, such as in the Florida mosquito populations used in this study, so that their actual role in permethrin detoxification can be assessed and possible resistance countermeasures implemented.

In summary, we identified the potential mechanisms involved in permethrin resistance in three *Aedes aegypti* populations from Florida. The p-s population had higher homozygous mutations in S989P, V1016I and F1534C that correlated to permethrin resistance level. The results also revealed two P450 genes, *CYP6N12* and *CYP6M11*, might be the potential enzymes that metabolize permethrin as it was induced after permethrin exposure. Further experiments need to be completed to identify the roles *CYP6N12* and *CYP6M11* play in permethrin resistance. Taken together, the results suggest that high permethrin resistance may be caused by several factors, including point mutations and xenobiotic metabolism. The effect of high resistance on adult longevity might change vectorial capacity which would increase the risk of mosquito-borne disease transmission.

---

**S1 Table.**
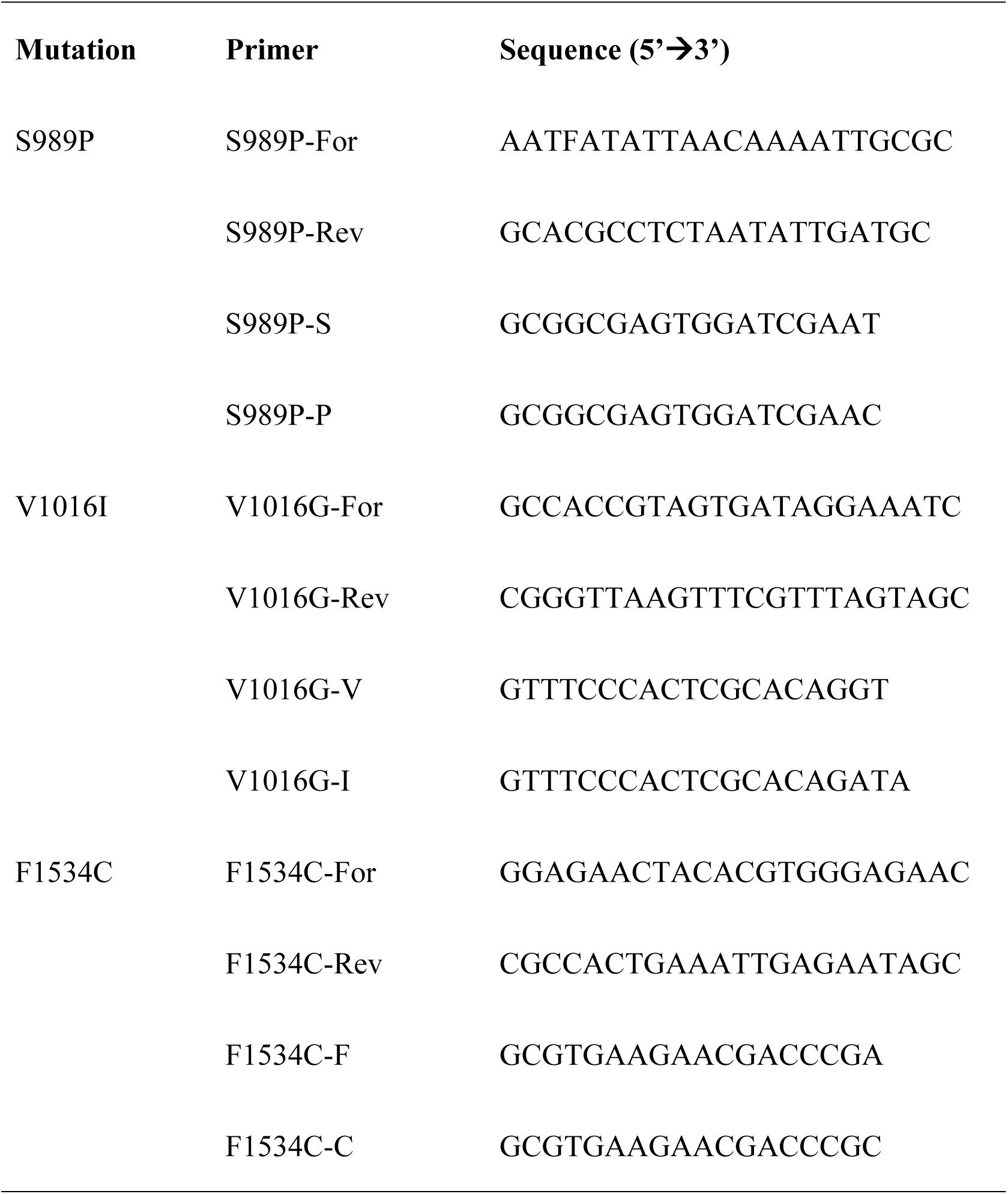
Primers used for the sodium channel gene mutation analysis.

**S2 Table.**
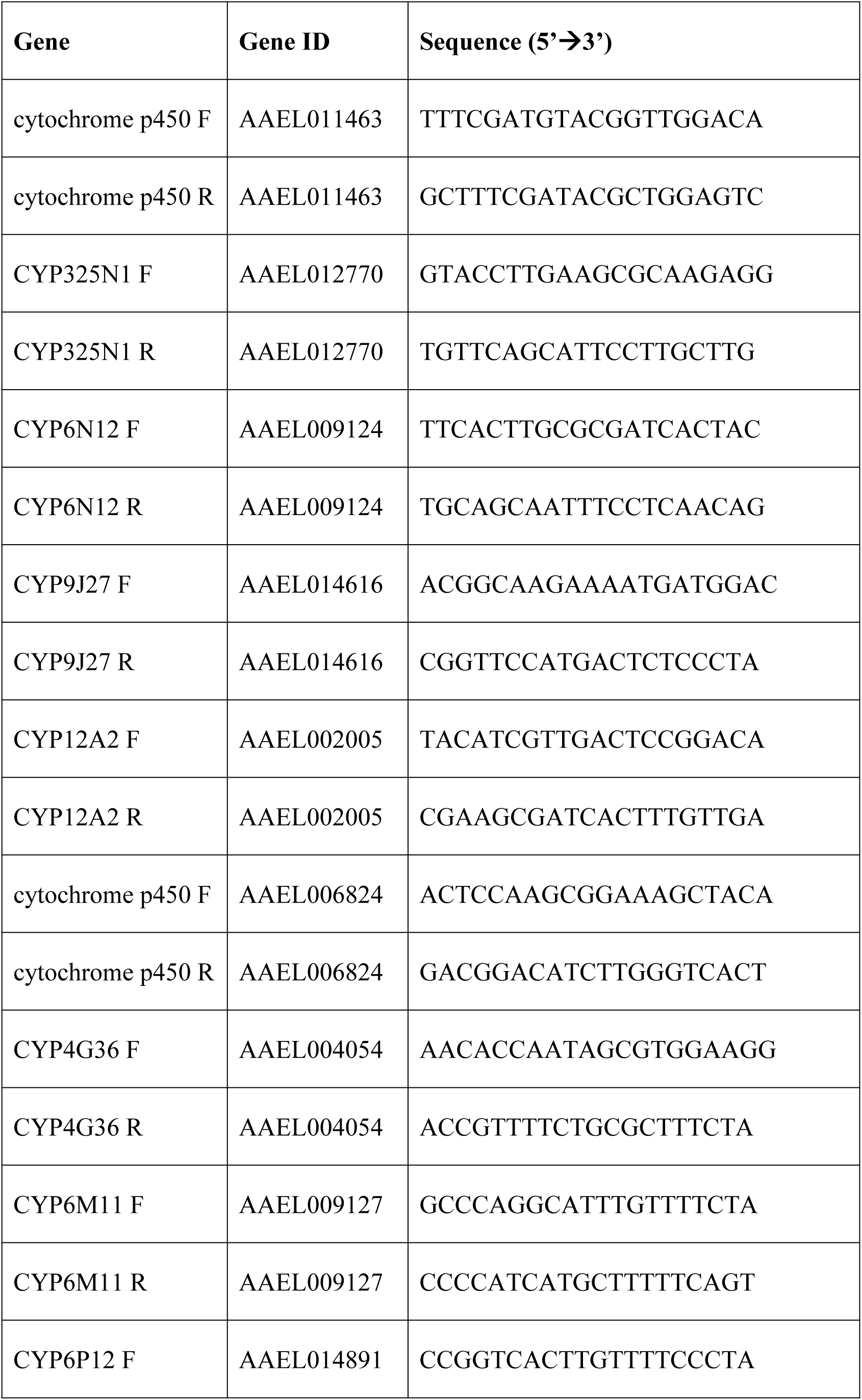
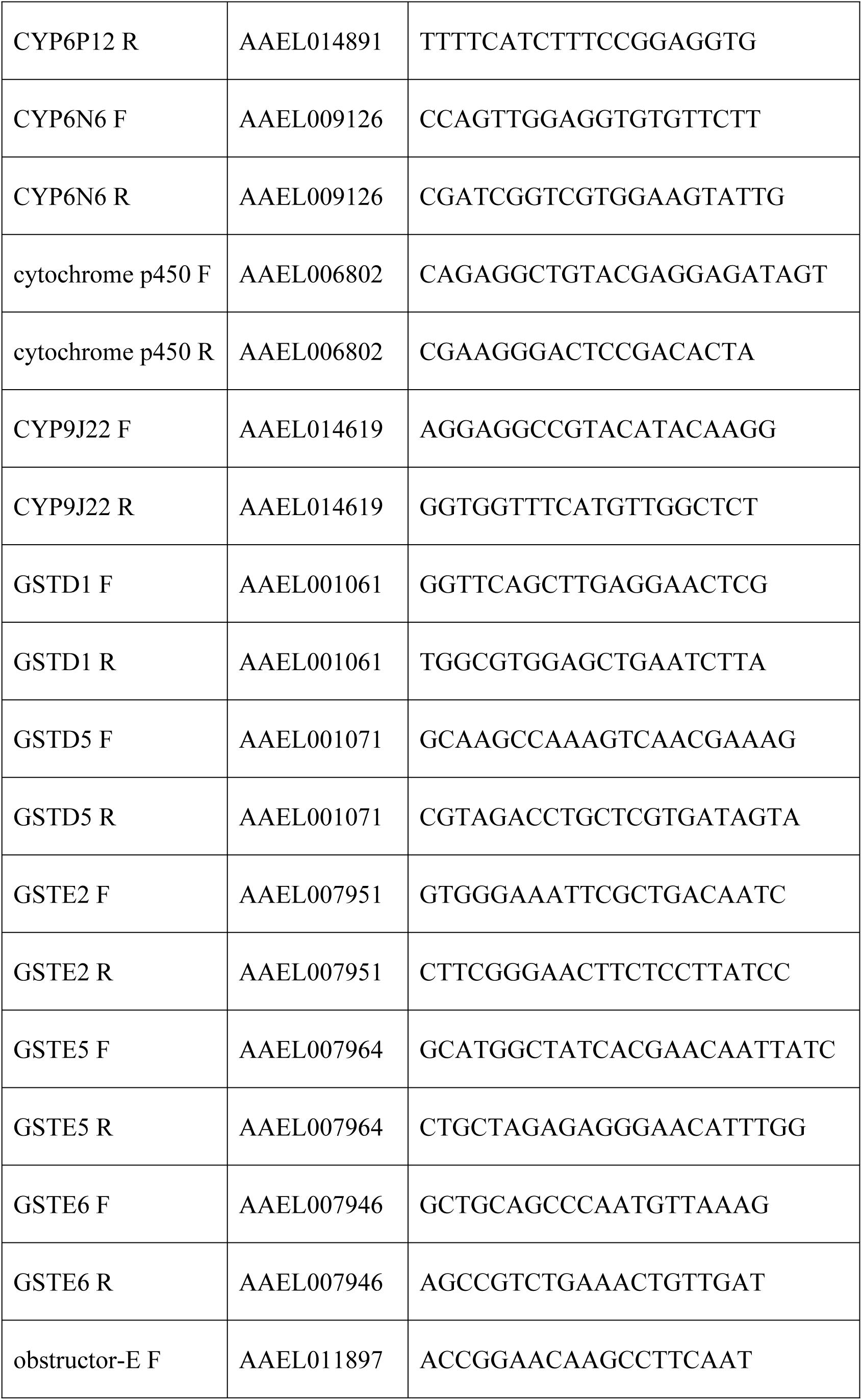
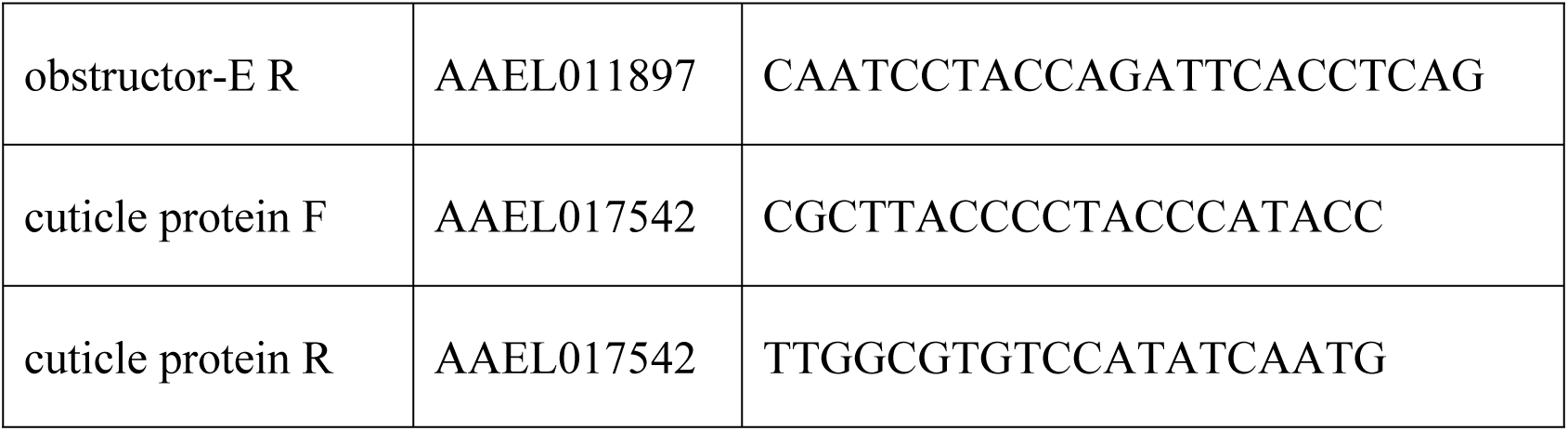
Primer sequences for the insecticide detoxification genes.

**S1 Figure.**
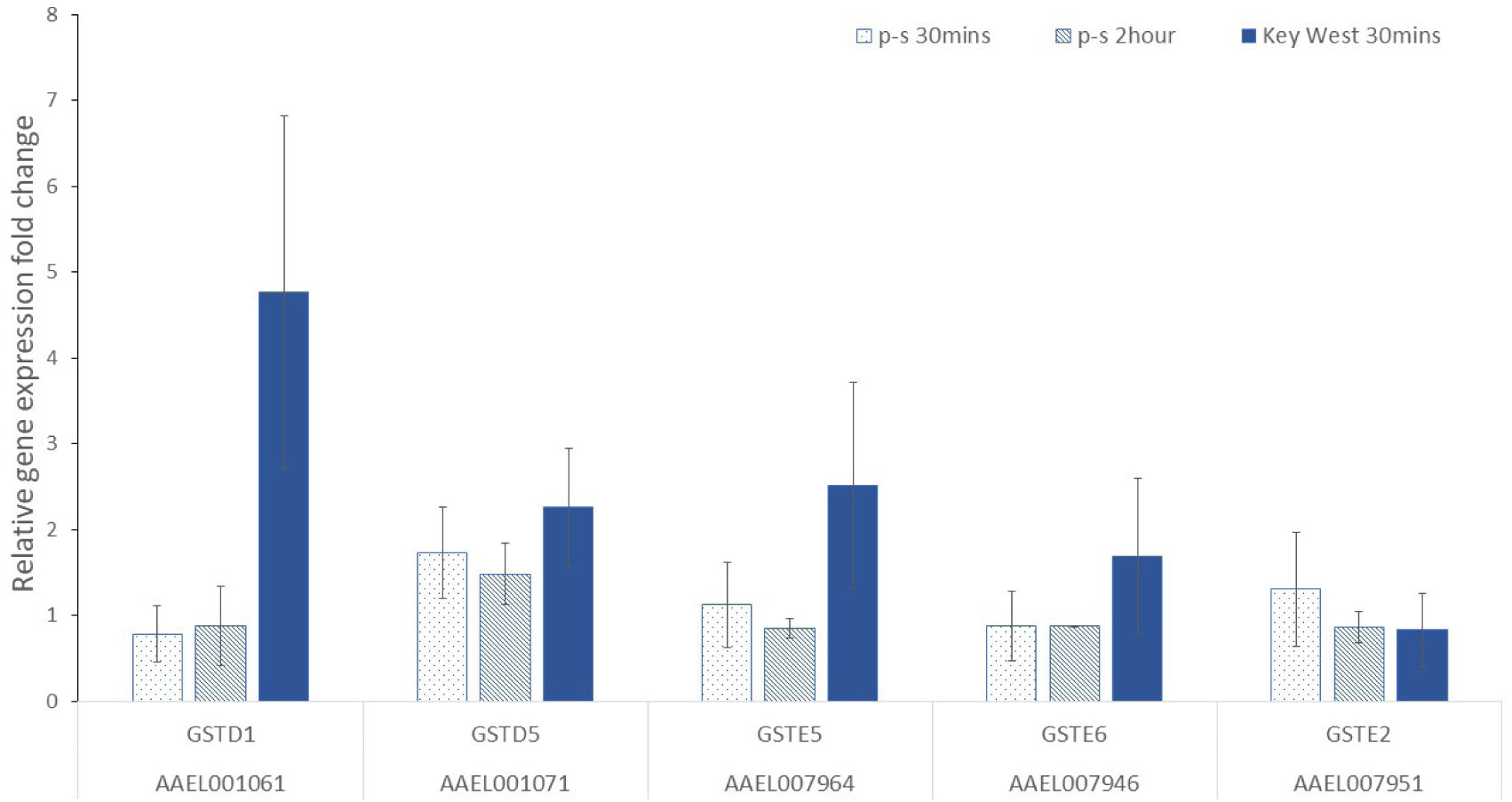
Comparison of glutathione-S transferase gene expression after permethrin treatment in the p-s population and the parental population (Key West) compared to non-treatment group.

